# Four cell wall-degrading enzymes of *Xanthomonas campestris* pv. *campestris* determine bacterial escape from hydathodes to the leaf vasculature

**DOI:** 10.1101/2024.10.09.617435

**Authors:** Misha Paauw, Marcel Giesbers, Sebastian Pfeilmeier, Harrold A. van den Burg

**Affiliations:** Molecular Plant Pathology, Swammerdam Institute for Life Sciences (SILS), Faculty of Science, University of Amsterdam, Science Park 904, 1098 XH, Amsterdam, the Netherlands; Wageningen Electron Microscopy Centre, Department of Plant Sciences, Wageningen University & Research, Droevendaalsesteeg 4, 6708 PB, Wageningen, the Netherlands

**Keywords:** Type II secretion system, tissue-specificity, cell wall degrading enzymes, vascular colonization, *Xanthomons campestris*, Black rot disease

## Abstract

To colonize plants, pathogenic bacteria modulate the biology of the host employing different bacterial secretion systems. For example, the type II secretion system (T2SS) releases toxins, proteases, lipases and carbohydrate-degrading enzymes into the extracellular environment to promote tissue softening and soft rot. In this way, the T2SS promotes virulence of phytopathogenic Gram-negative bacteria. However, the role of the T2SS and its substrates for vascular disease remains enigmatic. Here, we show that the Xps-T2SS allows *Xanthomonas campestris* pv. *campestris* (Xcc) to breach the tissue barrier between hydathodes– the initial bacterial entry point – and xylem thereby gaining access to the leaf vasculature. Yet, Xps-T2SS was dispensable for bacterial multiplication in the leaf apoplast or inside the hydathode cavity suggesting a role beyond plant defense suppression or nutrient acquisition. Using comparative genomics, four plant cell wall-degrading enzymes (CWDEs) were found to be associated with vascular pathogenesis. Testing gene knockout combinations of those enzymes revealed that virulence of only the quadruple CWDE mutant was down to the level of the *xps*-T2SS mutant. Our results thus demonstrate that the Xps-T2SS and a set of CWDEs that is likely secreted by this system allow Xcc to break this tissue barrier enabling long-distance mobility of Xcc inside the host plant. We thus expand our understanding on how certain bacterial pathogens have specialized towards vascular pathogens.

## Introduction

The general notion that most plants are resistant to most pathogens underlines the effective defence mechanisms of plants against invading microbes, and the sophisticated virulence mechanisms that pathogenic microbes have evolved to overcome these defence mechanisms (Panstruga & Moscou, 2020; Zhang *et al*., 2020). Successful pathogens enter plant tissue and establish a suitable micro-environment for proliferation by suppressing plant immune responses, acquiring nutrients and creating an aqueous microhabitat (Roussin-Léveillée *et al*., 2024). To achieve this, bacterial pathogens employ different protein secretion systems to actively manipulate and combat a variety of host processes and barriers (Alvarez-Martinez *et al*., 2021; Cianciotto & White, 2017; Coombes, 2009). For example, the type III secretion system (T3SS), which is required for bacterial virulence, has evolved to inject type III secreted effectors (T3Es) into the host cytosol to manipulate intracellular processes (Roussin-Léveillée *et al*., 2024; Schreiber *et al*., 2021). In contrast, the type II secretion system (T2SS) releases proteins into the extracellular environment, including enzymes that degrade plant cell walls, of which the breakdown products presumably serve as bacterial nutrients (Alvarez-Martinez *et al*., 2021; Cianciotto & White, 2017; Solé *et al*., 2015). Although the T2SS is not an exclusive feature of pathogenic bacteria (Cianciotto, 2005; Cianciotto & White, 2017; Jha *et al*., 2005), this secretion system is required for full pathogenicity in a wide range of pathogens of both plants and animals, including *Xanthomonas oryzae* (Ray *et al*., 2000), *X. campestris* (Dow *et al*., 1987; Hu *et al*., 1992; Qian *et al*., 2005), *X. vesicatoria* (Szczesny *et al*., 2010), *Erwinia carotovora* (Pirhonen *et al*., 1991), *Dickeya* spp. (Condemine & Le Derout, 2022; Ji *et al*., 1989) *Ralstonia solanacearum* (Tsujimoto *et al*., 2008), *Xylella fastodiosa* (Ingel *et al*., 2023), *Burkholderia gladioli* (Chowdhury & Heinemann, 2006), *Legionella pneumophila* (Rossier *et al*., 2004), and *Citrobacter rodentium* (Krekhno *et al*., 2024). However, it remains poorly understood which step of the infection process requires T2SS.

Plant pathogenic bacteria of the genus *Xanthomonas* are not only phylogenetically grouped in species. Certain species are further subdivided in host-specific clades called pathovars (pv). Importantly, pathovars of different *Xanthomonas* species also form two distinct groups based on their tissue-specificity on their respective host, that is, they cause (a) vascular infections or (b) colonize the extracellular space (apoplast) of the leaf mesophyll (Gluck-Thaler *et al*., 2020). For example, *X. campestris* pv. *campestris* (Xcc) invades plants belonging to the Brassicacea family via hydathodes, organs at the leaf margin, prior to gaining access to the leaf vasculature (Cerutti *et al*., 2017; Paauw *et al*., 2023). To do this, Xcc causes degradation of the primary cell walls of xylem tracheary endings that irrigate the hydathode epithem tissue (Bretschneider *et al*., 1989; Cerutti *et al*., 2017). In contrast, the stomata-invading *X. campestris* pv. *raphani* (Xcr) encounters a different host environment when colonizing the apoplast via stomatal openings. Despite being closely related with an average nucleotide identity (ANI) >97% across the genomes (Paauw *et al*., 2024b), both pathogens have adapted to thrive in their respective niche on the same host plant. In line with the observation of primary cell wall degradation in xylem tissue by Xcc, the cell wall-degrading enzyme (CWDE) CbsA was found to be associated with vascular pathogenicity in the genera *Xanthomonas* and *Xylella* (Gluck-Thaler *et al*., 2020). CbsA is a 1,4-β-cellobiohydrolase (EC 3.2.1.19, glycosylhydrolase family GH6) that hydrolyzes the non-reducing ends of cellulose chains, thereby releasing the disaccharide cellobiose (Tayi *et al*., 2018). For the vascular pathogens *X. oryzae, R. solanacearum,* and *X. fastodiosa*, CbsA is required for full vascular pathogenicity (Gluck-Thaler *et al*., 2020; Jha *et al*., 2007; Liu *et al*., 2005) and heterologous expression of CbsA in otherwise non-vascular pathovars of *X. translucens* is sufficient to confer vascular pathogenicity (Gluck-Thaler *et al*., 2020). However, in other *Xanthomonas* species, such as Xcc, the impact of deleting the *cbsA* gene has not yet been reported and the transfer of *cbsA* from Xcc to the non-vascular Xcr did not alter tissue-specificity of Xcr (Dubrow *et al*., 2022). Therefore, it was hypothesized that additional factors must contribute to vascular pathogenicity of Xcc, but these factors remained elusive so far (Dubrow *et al*., 2022).

Here, we generated T2SS and T3SS mutants in a bioluminescent reporter strain of Xcc (Paauw *et al*., 2023; van Hulten *et al*., 2019). The reporter strain allowed us to study Xcc spread within plant leaves with a high spatiotemporal resolution following a natural infection—from hydathode colonization to the vasculature and finally into the leaf mesophyll (Paauw *et al*., 2023). Although both T2SS and T3SS mutants did not cause visible disease symptoms, we reveal striking differences in leaf colonization patterns between T2SS and T3SS mutants, highlighting a role for the T2SS in Xcc to access and to break out from the vasculature. To identify niche-specific adaptations, a pangenome comparison was made between isolates of the vascular Xcc and non-vascular Xcr. We discovered a set of four genes encoding potential T2SS substrates that combined are critical for vascular pathogenesis.

## Results

### Xps*-*type II secretion system is required for tissue-specific pathogenicity in Xcc

The Xcc genome encodes two T2SS gene clusters: (i) the *xps-*T2SS cluster, which is required for pathogenicity (Dow *et al*., 1987; Hu *et al*., 1992; Qian *et al*., 2005) and is conserved throughout the genus *Xanthomonas* (Alvarez-Martinez *et al*., 2021), and (ii) the *xcs*-T2SS cluster. However, it is unclear whether the requirement of Xps*-*T2SS for pathogenicity is (i) tissue-specific and (ii) involved in breaching the tissue barrier from the hydathode to the xylem. To examine both, we generated single and double loss-of-function mutants of the Xps- and Xcs-T2SS by deleting the key genes *xpsD* and/or *xcsD*, similar to earlier studies on Xcc and *Xanthomonas campestris* pv. *vesicatoria* (Chen *et al*., 1996; Szczesny *et al*., 2010). As genetic background, we used a bioluminescent reporter strain of Xcc8004 that lacks the type III effector XopAC (AvrAC) to avoid activation of the ZAR1-mediated immune response in *Arabidopsis thaliana* (hereafter Arabidopsis), similar to previous studies (Paauw *et al*., 2023; Wang *et al*., 2015). This bacterial strain is henceforth referred to as ‘wildtype’ (WT) and is highly virulent on the susceptible Arabidopsis accession Oy-0 (Paauw *et al*., 2023). To study colonization of the hydathode, midvein, and mesophyll, we performed spray, clipping and syringe inoculations on this accession, respectively. Following spray inoculation, both the *ΔxpsD* single and *ΔxpsD*/*ΔxcsD* double mutants failed to spread from infected hydathodes (at 7 days post inoculation [dpi]) towards the leaf vasculature and mesophyll at 14 dpi, unlike WT and the *ΔxcsD* single mutant. (Fig 1A and 1B, Fig S1). Importantly, the *ΔxpsD* single mutant and the double mutant *ΔxpsD/ΔxcsD* were both not impaired hydathode colonization visualized at 7 dpi (Fig 1A and Fig 1C). Furthermore, the *ΔxpsD* mutant was still metabolically active and it formed high density bacterial colonies in the hydathodes of the majority of the examined plants at 14 dpi (luminescence index = 1; Fig 1D). In contrast, and as expected, the T3SS mutant *ΔhrcV* seldomly reached bacterial densities resulting in a detectable bioluminescence signal in hydathodes and it was entirely incapable of causing a systemic infection along the leaf vasculature (Fig 1). Taken together, these observations suggest that the *ΔxpsD* mutant is capable of entering the hydathode cavity via guttation droplets and subsequently proliferate to a high sustainable bacterial density in the cavity. Yet, *ΔxpsD* was incapable of reaching the leaf vasculature of Arabidopsis, the next step in a natural infection.

**Figure 1.**
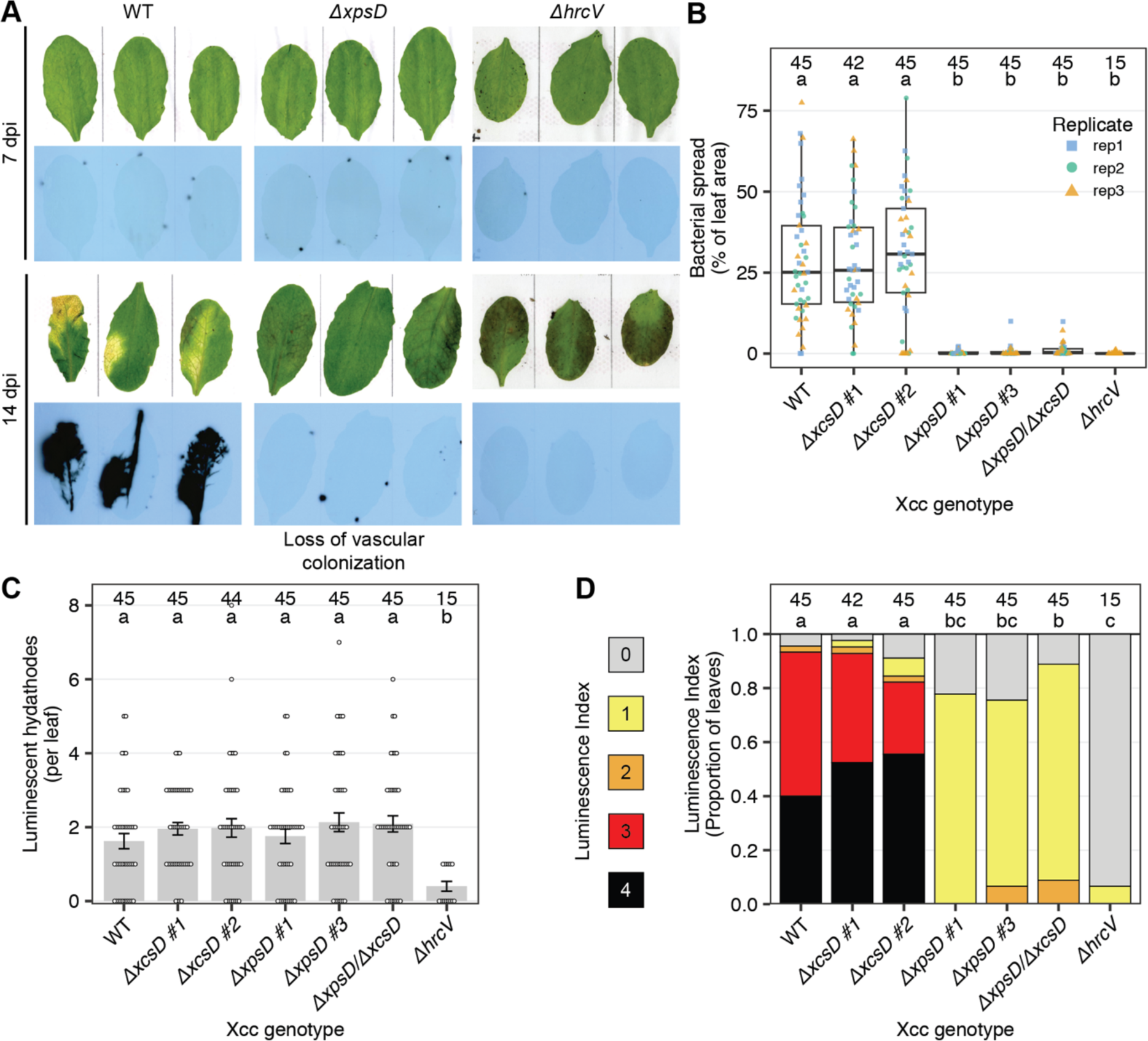
*xps-*T2SS is required for colonization of the leaf vasculature upon hydathode entry. Arabidopsis Oy-0 plants were spray inoculated with Xcc *ΔxopAC Tn*7*:lux:mTq2* (WT) and indicated mutant strains. (A) Representative images of Oy-0 leaves inoculated with Xcc WT and selected mutants. For both 7 and 14 dpi, top rows show scans of infected plant leaves, while the bottom row shows bacterial luminescence in the same leaves. Additional representative images of other mutants are shown in Figure S1. (B) Quantification of bacterial spread at 14 dpi, measured as ‘% of the total leaf area’. Individual data points are shown as squares, circles, or triangles depending on the experimental replicate. (C) Quantification of the number of luminescent hydathodes per leaf at 7 dpi. Individual data points are shown as dots. Error bars represent the standard error of the mean. (D) Quantification of bacterial spread at 14 dpi by using ordinal luminescence indices. In panel B-D, three independent experiments were combined and the total sample size is indicated above the bars or boxplots. Groups with no letter in common have a significantly different mean or mean rank (Tukey’s or Dunn’s; α = 0.05).

Next, we tested whether the *ΔxpsD* mutant was still able to proliferate and to display long distance movement along the leaf vasculature when the mutant was directly introduced into the vasculature thereby bypassing the natural tissue barrier between hydathodes and the vasculature. To do this, we performed clipping assays removing the leaf tip and quantified both (a) the distance along the midvein colonized by the bacteria and (b) the total leaf area colonized (Fig 2A). Remarkably, the *ΔxpsD* mutants were still able to spread along the entire length of the leaf midvein in these clipping assays, albeit slower than WT and *ΔxcsD* (Fig 2B and 2C). In contrast, the T3SS mutant *ΔhrcV* apparently failed to move away from the initial wound site (Fig 2B and 2C). Despite the capacity of the *ΔxpsD* to colonize the entire midvein, both the total leaf area colonized and the symptom development (chlorosis) were still reduced for *ΔxpsD* compared to WT and *ΔxcsD* (Fig 2B and 2D). In corroboration, we did not observe *ΔxpsD* spreading from the midvein into lateral veins or the adjacent mesophyll tissue, highlighting a potential role for the Xps-T2SS in spread of Xcc towards the lateral veins and apoplast.

**Figure 2.**
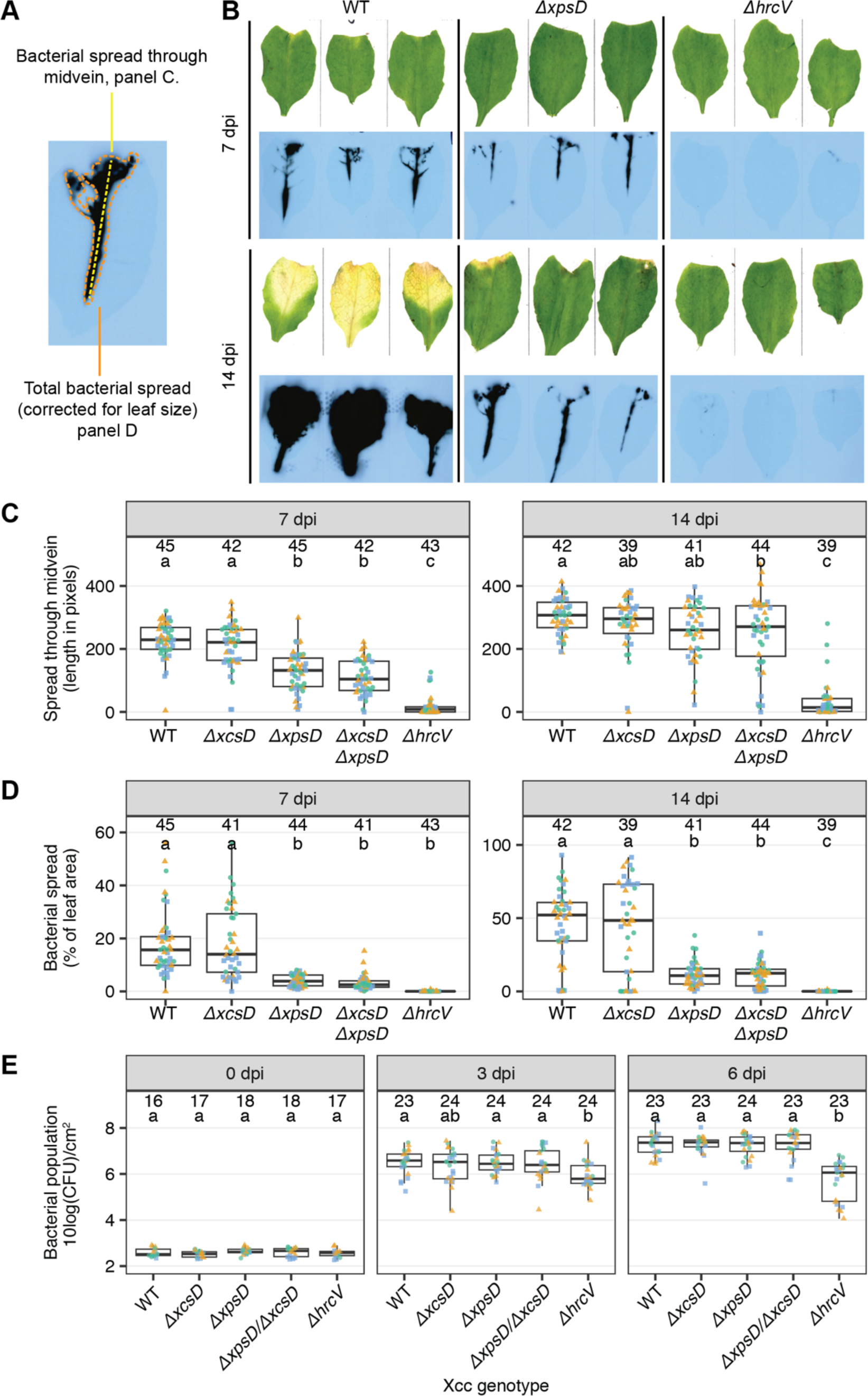
The Xps type II secretion system is required for bacterial spread from the midvein to lateral veins, while it is dispensable for multiplication in the apoplast. (A) Illustration of bacterial spread quantification shown in panels C and D. (B) Representative images of Arabidopsis Oy-0 leaves after leaf clip inoculation of Xcc *ΔxopAC Tn*7*:lux:mTq2* (WT) or indicated mutant lines. (C) Quantification of bacterial spread through the midvein of Arabidopsis leaves at 7 and 14 days after clip inoculation. (D) Quantification of bacterial spread over entire Arabidopsis leaves as ‘% of the total leaf area’ at 7 and 14 days after clip inoculation. (E) Bacterial population size directly after (0 days post inoculation; dpi), three, and six days post syringe infiltration into the leaf apoplast. In panels C, D and E, data from three independent experiments was combined resulting in a total sample size indicated above the boxplots, and individual data points are colored by the independent experimental replicate. Groups with no letter in common have a significantly different mean (panel E) or mean rank (panel C and D) (Tukey’s or Dunn’s; α = 0.05).

Finally, to assess whether the T2SS mutants showed a difference in bacterial multiplication in the leaf apoplast, growth of the bacterial population was determined in the apoplast following syringe inoculations. The T2SS mutants multiplied to a similar population size as the WT at three and six days post inoculation (dpi) (Fig 2E). In contrast, the negative control *ΔhrcV* reached approximately a three magnitudes lower population density at six dpi. Taken together, these findings imply that Xcc requires the Xps*-*T2SS during a natural infection cycle, in particular for crossing different plant tissue barriers, thereby enabling entry into the vasculature and escape from the vasculature towards the mesophyll.

### *ΔxpsD* forms biofilms inside xylem vessels

To confirm that *ΔxpsD* mutants were able to proliferate inside xylem vessels of plant leaves, we made longitudinal cross sections of the leaf vasculature of *Brassica oleracea* plants and inspected these leaf cross sections using scanning electron microscopy (SEM). First, we studied non-inoculated leaf material and focused on the junctions between the midvein and side veins (Fig S2A and S2B). We observed long xylem vessels (tracheids) lined with spiraling secondary cell wall thickenings. In the vertical direction of xylem vessels, no cell wall barriers were observed that could obstruct microbial colonization, while the side walls of a vessel showed numerous pits, which appeared to be filled with a porous structure and connected either to a neighboring vessel or the intercellular space (Fig S2C), as described in the literature (Kashyap *et al*., 2021; Yadeta & Thomma, 2013). Upon inoculation with both WT and *ΔxpsD*, we detected a thick bacterial biofilm that covered the entire cell wall of the xylem tracheids in the infected leaves (Fig S2D).

### Four cell wall-degrading enzymes are conserved in Xcc and absent in non-vascular pathovar Xcr

The T2SSs secretes proteins into the surrounding extracellular space and includes a cocktail of CWDEs (Alvarez-Martinez *et al*., 2021; Solé *et al*., 2015). Therefore, we hypothesized that Xcc relies on its T2SSs to secrete a specific repertoire of CWDEs that enables Xcc to switch towards a vascular lifestyle. To identify these CWDEs, we performed a pangenome analysis of Xcc and Xcr using a recently described dataset of whole genome assemblies from a diverse collection of *Xanthomonas campestris* isolates, both in year of isolation and region of isolation (Paauw *et al*., 2024b). In total, we predicted for 49 Xcc and 44 Xcr genomes all the open reading frames (ORFs) followed by a genome annotation and clustering of the annotated ORFs (genes) into orthogroups using Roary (Page *et al*., 2015) (Fig 3A). This resulted in a total of 11,824 orthogroups, of which the majority is present (a) in only single individual isolate or (b) conserved across all genomes (Fig S3A). In addition, our isolate collection contains a representative pangenome as rarefaction curves tail off indicating that additional genomes would add few novel orthogroups (Fig S3B). Consistent with our hypothesis that there is an orthogroup-based difference between the Xcc and Xcr genomes, the genomes from the different pathovars clustered separately in a multiple correspondence analysis (Fig 3B).

**Figure 3.**
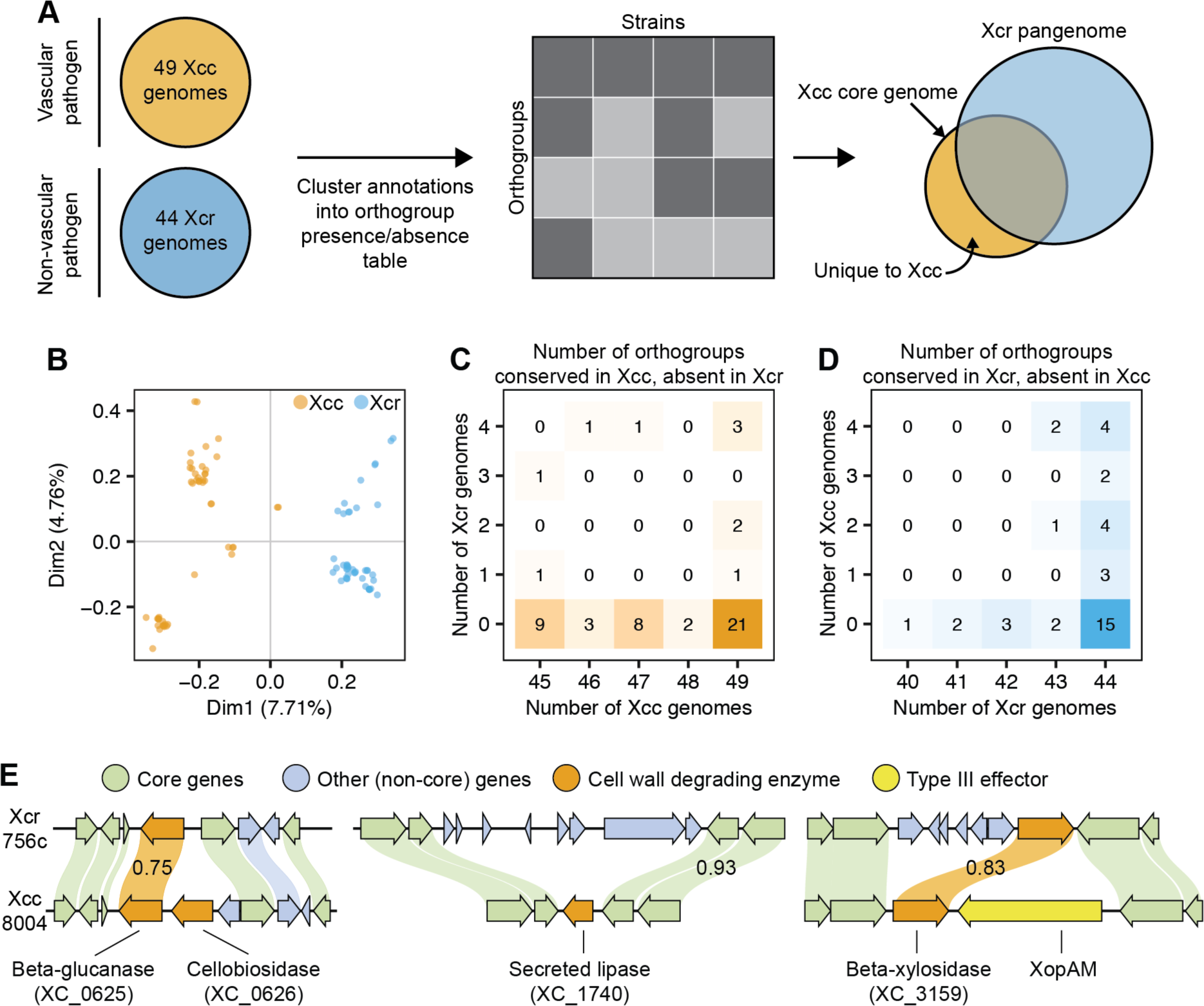
Four plant CWDEs are present in (nearly) all Xcc genomes while absent in all Xcr genomes. (A) Overview of the pangenome pipeline used to identify Xcc-specific orthogroups. (B) Multiple correspondence analysis based on an orthogroup presence/absence matrix. (C) Number of orthogroups present in *n* Xcc genomes (x-axis) and *m* Xcr genomes (y-axis), thus highlighting orthogroups conserved among Xcc and absent in Xcr. (D) As E, except that this plot focusses on orthogroups conserved in Xcr and absent in Xcc. (E). Comparison of the genomic loci encoding Xcc-specific CWDEs with the corresponding locus in Xcr (determined by the core genes around the Xcc-specific CWDEs). Arrows represent open reading frames. Vertical links between ORFs represent orthologous relationships between ORFs with sequence identity > 0.95, unless otherwise noted in the link.

We then looked for orthogroups that divide the two pathovars into two separate groups, i.e. orthogroups present in (nearly) all genomes of one pathovar while absent in the other (Fig 3C and 3D). Using these criteria we detected 31 orthogroups unique for Xcc (with the conservation threshold set at ‘present in at least 47 out of the 49 Xcc genomes analyzed’) (Table S1). This set included four orthogroups annotated as CWDE (hereafter: Xcc-specific CWDEs). In specific, we detected two neighboring genes encoding a cellulase (CAZy family: GH5_1, gene identifier: XC_0625) and a 1,4-β-cellobiohydrolase CbsA (GH6, XC_0626) previously associated with vascular pathogenicity in the genus *Xanthomonas* (Gluck-Thaler et al., 2020). In addition, we detected a gene encoding the secreted lipase LipA (XC_1740), of which homologs in other *Xanthomonas* and *Xylella* species were shown to contribute to virulence and the *Xyllella fastidiosa* homolog was shown to be T2SS secreted (Aparna *et al*., 2009; Nascimento *et al*., 2016; Tamir-Ariel *et al*., 2012). Furthermore, we detected a gene annotated as β-xylosidase (GH3, XC_3159). Of these four genes, XC_0626 and XC_1740 were absent in the corresponding genomic position in Xcr. In the case of XC_0625 and XC_3159 a homologous gene was present at the corresponding genomic position in Xcr, but the sequence similarity was below the orthogroup threshold (Fig 3E). Indeed, the Xcr homologs of these two CWDEs were found in the set of 20 Xcr-specific orthogroups (Fig 3D and Table S2). Besides CWDEs, we found five Xcc-specific T3Es, that is, AvrBs1, XopAG, XopAY, XopQ, and XopAM as reported before (Paauw *et al*., 2024b). XopAM flanks the Xcc-specific β-xylosidase described above (Fig 3E). Finally, we detected *fliC*, which encodes the main building block of the bacterial flagellum, and the *Ef-Tu* gene encoding Elongation factor Tu as orthogroups specific to both Xcc and Xcr, confirming pathovar-specific protein sequence for these two plant immunogenic molecules (Roux *et al*., 2015). Taken together, our pangenome analysis revealed four Xcc-specific CWDEs that could contribute to the specialization of Xcc to a vascular lifestyle of a bacterial plant pathogen.

### Xcc-specific CWDEs contribute to vascular colonization by Xcc

To assess the contribution of the Xcc-specific CWDEs to vascular disease, we generated the corresponding gene deletion mutants of these enzymes in our bioluminescent WT strain (Xcc8004 *ΔxopAC Tn*7*:lux:mTq2*). In addition, we stacked the gene deletions to reveal any additive, synergistic, or complementary effects of the encoded enzymes on Xcc virulence and Black rot disease symptom development. We found that, for the single gene mutants, only the *cbsA* mutant *ΔXC_0626* showed reduced vascular colonization, but not to the same degree as the T2SS double mutant *ΔxpsD/ΔxcsD* (Fig 4A). None of the three double mutants, nor two triple mutants (all in addition to the *ΔXC_0626* mutation) showed an additional reduction in bacterial spread (Fig 4A and 4B). In contrast, the quadruple mutant lacking all four Xcc-specific CWDEs (*ΔXC_0625*/*ΔXC_0626/ΔXC_1740/ΔXC_3159)* displayed a strong reduction in bacterial virulence similar to *ΔxpsD/ΔxcsD* in our natural infection assay (Fig 4C). Yet, in contrast to the T2SS mutant *ΔxpsD,* the quadruple CWDE mutant was still able to degrade leaf material (Fig S4), as tested on leaf discs of the Arabidopsis mutant *rbohD* (Pfeilmeier *et al*., 2024). This suggests that (i) the four Xcc-specific CWDEs are not critical to degrade leaf mesophyll tissue, instead, they are apparently specialized to vascular pathogenesis and (ii) secretion of other lytic enzymes is not impaired in the quadruple CWDE mutant. In addition, the quadruple CWDE mutant displayed similar growth curves as wildtype in both rich and minimal liquid medium (Fig S5). Finally, complementation of *ΔXC_0626* by reintroducing the XC_0626 coding sequence into the genome of the Xcc *ΔXC_0626* mutant could restore bacterial virulence to WT levels (Fig 4C). Taken together, these experiments suggest that the quartet of Xcc-specific CWDEs together contribute to vascular spread of Xcc in Arabidopsis leaves, with the largest contribution by CbsA (XC_0626).

**Figure 4.**
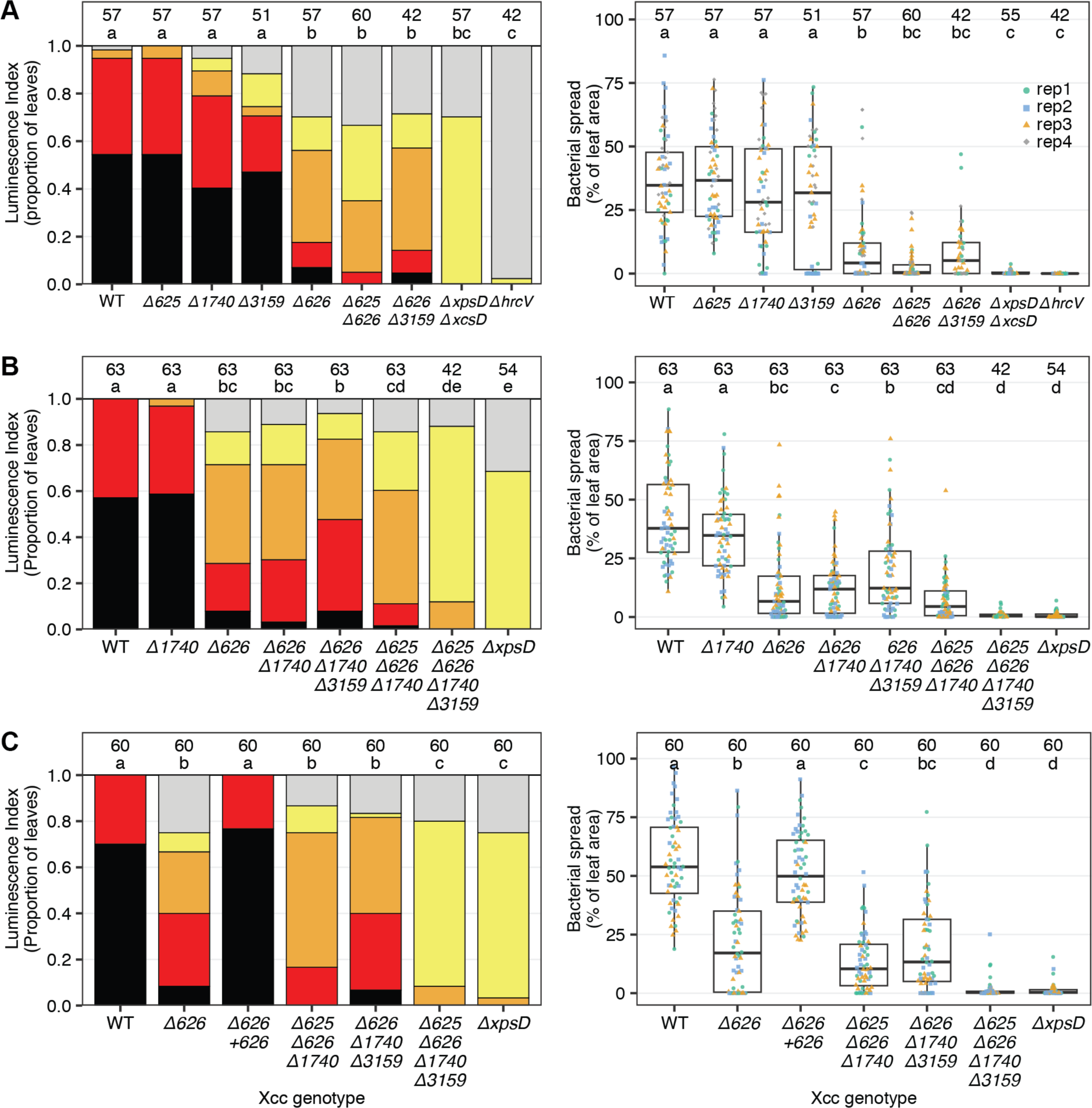
Four Xcc-specific CWDEs together contribute to vascular colonization by Xcc upon hydathode colonization, with the largest contribution for CbsA. Arabidopsis Oy-0 plants were spray inoculated with Xcc *ΔxopAC Tn*7*:lux:mTq2* (WT) and indicated CWDE mutants in this background. The CWDE mutants are indicated with a delta sign followed by the number of their gene identifier: *Δ625*: β-glucanase (XC_0625), *Δ626*: cellobiosidase (XC_0626), *Δ1740*: secreted lipase (XC_1740), *Δ3159*: β-xylosidase (XC_3159). In each panel, two graphs display the bacterial spread in Arabidopsis leaves quantified with an ordinal luminescence index (left) and the percentage of leaf area (right). For each panel, data from three or four independent experiments were combined. The total sample size is indicated above the bars or boxplots and individual data points are colored by the independent experimental replicate. Groups with no letter in common have a significantly different mean rank (luminescence index) or mean (bacterial spread) (Dunn’s or Tukey’s α = 0.05, respectively).

## Discussion

In this study, we investigated the evolutionary adaptations of Xcc to the vascular environment of plants in comparison to Xcr. These adaptations revolve around the key question of how the T2SSs contribute to bacterial spread from infected hydathodes into xylem endings and afterwards systemically into the vasculature. To investigate this, we tracked leaf colonization of Xcc T2SS mutants with high spatiotemporal resolution using our bioluminescence reporter system (Paauw *et al*., 2023; van Hulten *et al*., 2019). We demonstrated that *xps-*T2SS and not *xcps-*T2SS mutants are strongly compromised in (i) reaching the vasculature upon natural infection via hydathodes and (ii) movement towards lateral veins and the leaf mesophyll upon clipping inoculations, which provides direct access to the midvein. Yet, the *xps-*T2SS mutant bacteria were still able to (i) colonize hydathodes, (ii) spread through the entire leaf midvein, and (iii) multiply to WT levels in the leaf apoplast, in contrast to the T3SS mutant *ΔhrcV*. Indeed, our results show that the T3SS is required for both hydathode colonization and colonization of the midvein, and is required for full virulence in the mesophyll. The striking difference in leaf colonization between the *xps-*T2SS and T3SS loss-of-function mutants indicates that the T3SS is required for pathogenicity in general in all the tissues tested, while the T2SS seems to be important for specific steps of the Xcc infection cycle enabling long-distance movement of Xcc. These observations go beyond previous studies, as both T2SS and T3SS mutants cause no visible disease symptoms (Fig 1 and Fig 2), and until now, it was generally accepted that Xcc requires the Xps-T2SS for general pathogenicity just like the T3SS (Dow *et al*., 1987; Hu *et al*., 1992; Qian *et al*., 2005).

Our findings are in line with the hypothesis that bordered pits in the xylem vessels restrict microbial passage (Choat *et al*., 2008; Kashyap *et al*., 2021; Yadeta & Thomma, 2013). These pits contain a (hemi)cellulose and pectin membrane which is permeated by numerous pores that are too narrow for bacteria to pass, but allow xylem sap to flow freely (Choat *et al*., 2008). Conceivably, T2SS secreted CWDEs dissolve the pit membrane inside pit elements, enabling the bacteria to spread between xylem vessels. Using SEM, we were able to observe pits in xylem vessels of healthy *B. oleracea* leaves, but not in infected plant material due to a bacterial biofilm covering the entire wall of the xylem vessels (Fig S2). In a recent study on *X. fastidiosa*, it was found that this bacterium widens xylem pits in susceptible olive cultivars (Montilon *et al*., 2023), and there is evidence that pit membrane degradation depends on likely T2SS secreted endoglucanases (Ingel *et al*., 2019). Although beyond the scope of this work, it would be interesting to image hydathodes and xylem vessels of plant marker lines using confocal (Newman *et al*., 2003) or transmission electron microscopy (Cerutti *et al*., 2017; Ellis *et al*., 2010) to observe xylem pit integrity upon infection with Xcc WT and T2SS mutants.

We identified four vascular-pathogen specific CWDEs potentially secreted by the T2SS that could explain the loss of pathogenicity of the *xps-*T2SS mutant and give insights into bacterial adaptations to the hydathode and xylem environment. In particular, a *cbsA* mutant in Xcc showed reduced but not a complete loss of pathogenicity—similar to what has been observed in other species such as *X. oryzae* (Jha *et al*., 2007) and *R. solanacearum* (Liu *et al*., 2005). Previously, it was hypothesized that additional factors were implicated in vascular colonization by Xcc besides the CWDE CbsA (Dubrow *et al*., 2022; Gluck-Thaler *et al*., 2020). Our pangenome analysis of a comprehensive collection of Xcc and Xcr isolates (Paauw *et al*., 2024b) revealed three additional CWDEs besides *CbsA* are specific for Xcc. A mutant strain lacking all four Xcc-specific CWDEs was strongly compromised in its pathogenicity to the extent seen for the *xps*-T2SS mutants, implying that these four CWDEs combined are required for vascular pathogenicity in Xcc. Two of these genes, CbsA and the secreted lipase, lacked a homolog in Xcr at the corresponding genomic location, indicating that these genes were recently gained in a common ancestor of the Xcc clade or lost in a common ancestor of Xcr clade. Based on our finding that Xcc appears to have diverged from the more ancestral Xcr clade, the former notion is the more favorable hypothesis (Paauw *et al*., 2024b). However, the flanking genomic regions did not show any clear sequence signatures of a recent horizontal gene transfer (HGT) such as aberrant GC content or proximity to transposable elements, in contrast to several Xcc-specific T3Es (Paauw *et al*., 2024b). For the other two Xcc-specific CWDEs, the β-glucosidase and β-xylosidase, a homologous protein was encoded in the same genomic position in the Xcr genomes, but the sequence identity was below the set threshold (sequence similarity > 95%) for orthogroup detection. Conceivably, the Xcc and Xcr homologs of these CWDEs have acquired a different substrate specificity since the phylogenetic split between Xcc and Xcr. A similar situation was recently described for the bacterium *Dickeya chrysanthemi—*the causal agent of soft rot diseases*—*with two CWDEs sharing 83% sequence identity but they displayed different substrate preferences to degrade xylan, depending on the monosaccharide decorations on the xylan backbone (Yu *et al*., 2024). This substrate specificity could be traced back to a single amino acid change between the two xylanase variants. Likewise, structural characterization of xyloglucan processing enzymes in *Xanthomonas* revealed that a single amino acid change was detrimental for the catalytic activity of a xylanase (Vieira *et al*., 2021). Considering this, it would be interesting to examine the substrate specificities of the Xcc and Xcr protein variants of the CWDEs described in this study. Similarly, investigating the architecture of the hydathode-xylem barrier including the cell wall composition, which might be accessible for the enzymatic activity of Xcc-specific CWDEs, would further advance our understanding of plant resistance mechanisms to this pathogen. Interestingly, variations in type, area, and spatial distribution of intervessel connections could not explain differences in *X. fastidiosa* susceptibility in a panel of resistant and susceptible grapevine genotypes (Fanton *et al*., 2024). In contrast, the pit membranes of susceptible grapevine cultivar Chardonnay appeared to be enriched in fucosylated xyloglucans, compared to the moderately resistant cultivar Cabernet Sauvignon (Ingel *et al*., 2019). Conceivably, it is the specific chemical composition of the intervessel pit membranes, in combination with the CWDE repertoire of the pathogen, that determines the outcome of vascular infections by bacteria such as Xcc and *X. fastidiosa*.

In summary, our study demonstrates a crucial role of the T2SS and four Xcc-specific CWDEs to escape from hydathodes and access the vasculature during natural Xcc infections. This finding implies an adaptation of Xcc virulence factors for this step of the infection process, and suggests the existence of a physical barrier between hydathode and vasculature.

## Materials & methods

### Bacterial strains, culture conditions, and transformations

All bacterial strains used in this study are derivatives of the model strain Xcc8004 and are listed in Table S3. In-frame deletion mutants were obtained by double recombination with derivatives of the suicide plasmid pOGG2 (Schulze *et al*., 2012). DNA fragments of 600 bp in length upstream and downstream of the gene-of-interest were selected to generate a fusion construct *in silico*. To ensure that the deletion of genomic fragments did not disrupt operon transcription, the start codon and the final three codons of the open reading frames were included in between the two flanking regions resulting in the following fusion constructs: *flankA-Start-Final3-FlankB*. The fusion fragments were synthesized (GenScript Europe) together with flanking *BsaI* cloning sites for Golden Gate cloning and inserted into pOGG2 (Table S4). The obtained suicide plasmids were electroporated into competent Xcc cells as described (Wang *et al*., 2016) and bacterial recombinants were selected on MOKA (8 g·L^-1^ casamino acids, 4 g·L^-1^ yeast extract, 2.4 g·L^-1^ K_2_HPO_4_·3H_2_0, 0.3 g·L^-1^ MgSO_4_·7H_2_0, and 15 g·L^-1^ Daishin agar) agarose plates supplemented with spectinomycin (100 µg·mL^-1^). To select for the second recombination step, a small scoop of bacterial cells (i.e., just visible on a pipet tip) was dissolved in 1 mL liquid MOKA medium, and 70 µL was plated on MOKA agarose plates supplemented with approximately 0.33 M sucrose. Colonies from these plates were restreaked to MOKA plates with and without spectinomycin, and spectinomycin sensitive colonies were genotyped by PCR.

For in-locus complementation, the genomic fragment encoding *cbsA* (XC_0626) was amplified along with approximately 600 bp upstream and downstream of the gene from Xcc8004 genomic DNA using primers FP12109 (5’-AGGTCTCACGACaataactccacacctttcg-3’) and FP 12110 (5’-AGGTCTCAATGGatccgatccagcagttg-3’), and cloned into pOGG2 as described above. In contrast to the knock-out transformations, we used triparental mating (Figurski & Helinski, 1979) to introduce the pOGG2 knock-in construct into Xcc cells. The double recombination procedure was then carried out as described above.

### Plant cultivation

In this study the *A. thaliana* accession Oy-0 was used, as this accession is hypersusceptible to Xcc8004 *ΔxopAC* (Paauw *et al*., 2023). *A. thaliana* plants were grown as described (Paauw *et al*., 2023; van Hulten *et al*., 2019). Briefly, seeds were stratified in the dark at 4 °C on moist filter paper for 3 days and then sown in 40-cell trays on potting soil (3:17 parts perlite:soil, Hol80 zaaigrond Nr1, Jongkind Grond, The Netherlands). The trays were covered with a translucent dome for 5 days to increase the relative humidity. Plants were grown at 22 °C, 70% relative humidity with a short-day light regime (11 hours of light, 13 hours of darkness).

### Xcc inoculum for plant infections

Xcc inoculum was prepared as described (van Hulten *et al*., 2019). Briefly, bacterial cultures were grown from −80 °C glycerol stocks on KADO agarose plates (MOKA medium supplemented with 10 g·L^-1^ sucrose) containing appropriate antibiotics (rifampicin 25 µg·mL^-1^, gentamycin 10 µg·mL^-1^) for 2 days at 28 °C. Bacteria were scraped from solid medium with a disposable spreader and resuspended in 10 mM MgSO_4_. This bacterial solution was then washed by centrifugation (3,000*g*, 10 min), decanting the supernatant, and resuspending the pellet in fresh 10 mM MgSO_4_. For spray and clip inoculations, the inoculum was then diluted in 10 mM MgSO_4_ to approximately 1 x 10^8^ colony forming units (CFU) per mL (OD_600_ = 0.1) and supplemented with 0.0002% Silwett L-77. For syringe infiltrations, the inoculum was diluted to approximately 1 x 10^6^ CFU per mL (OD_600_ = 0.0001) without adding Silwett L-77.

### Spray inoculation assays

Four-week-old Arabidopsis plants were spray inoculated as described (Paauw *et al*., 2023; van Hulten *et al*., 2019). Briefly, approximately 1x2 mL of bacterial inoculum was sprayed onto trays containing 20 Arabidopsis plants using an airbrush spray-gun, in a flow cabinet. The trays were then placed in a growth cabinet (Microclima MC1000, Snijders Labs). To promote guttation droplets to form at hydathodes, allowing natural entry of Xcc, a specific temperature/humidity cycle was used for the first 48 hours (van Hulten *et al*., 2019).

### Leaf clipping inoculation

Four-and-a-half-week-old Arabidopsis plants were used for clip inoculations. Fine dissection scissors were immersed in bacterial inoculum suspension, and approximately 5 mm of the tip of four fully elongated leaves per rosette was clipped off. Between each leaf clipping, the scissors were submerged in the bacterial suspension. Like after the spray inoculated plants, the clip inoculated plants were placed in a growth cabinet (Microclima MC1000, Snijders Labs) and underwent the same temperature/humidity cycle.

### Detection of Xcc luminescence *in planta*

For the spray- and clip inoculations, leaves were sampled at 7 and 14 dpi. The sampling procedure and detection of bacterial luminescence was performed as described (Paauw *et al*., 2023; van Hulten *et al*., 2019). Briefly, the three most diseased leaves per plant were sampled. In absence of visible disease symptoms, we selected leaves of similar developmental stage. The leaves were glued onto a 30×40 cm paper sheet with a preprinted grid. This sheet was covered with a thin transparency sheet and scanned on a flatbed scanner. Then, a light-sensitive film was placed on top to capture the bacterial luminescence using an overnight exposure of approximately 16 hours. The following day, the light-sensitive film was developed to reveal the spread of the bacteria in the leaves. To quantify the bacterial spread within the leaves, we used both manual assessment into ordinal luminescence indices (Paauw *et al*., 2023; van Hulten *et al*., 2019), and automated detection of bacterial luminescence signal using the ScAnalyzer pipeline (Paauw *et al*., 2024a). In addition, we measured the length of the bacterial spread into the leaf midvein in ImageJ (in pixels) for clip inoculations.

### Syringe inoculation assays

Four-and-a-half-week-old *A. thaliana* plants were used for syringe inoculations. The day prior to the experiment, the plants were placed inside propagators, to boost the relative humidity to nearly 100 % (as assessed visually by the appearance of condensation droplets on the propagator lids). Three or four fully elongated leaves per plant were infiltrated with bacterial inoculum using a needleless syringe. Then, the plants were allowed to air-dry for approximately one hour. The plants were then covered with a translucent propagator lid for one more day. For the remaining days, the slits on the propagator lids were opened and the lids were placed slightly ajar onto the propagator base, resulting in a relative humidity of approximately 80 %. The plants were kept at 22 °C, with a short-day light regime (11 hours of light, 13 hours of darkness) in a growth chamber. Leaf tissue samples were taken at 0, 3, and 6 dpi. Before sampling, leaves were surface sterilized by submerging entire leaves in 70% ethanol for 15 seconds, followed by two washes in sterile Milli-Q water. Each sample consisted of two leaf discs (diameter 5 mm) punched from the surface-sterilized leaves, taking care not to select an area damaged by the infiltration procedure. The leaf discs were lysed in 500 µL 10 mM MgSO_4_ using a TissueLyser II bead mill grinder (Qiagen). Then, 10× dilution series were made in 10 mM MgSO_4_, and 10 μl droplets of a dilution series were plated as streaks on KADO agarose plates with appropriate antibiotics. After two days of incubation at 28 °C, the colonies formed were counted in the dilution in which we could reliably count between 15 and 120 colonies.

### Pangenome analysis

We used the genome assemblies and annotations of Xcc and Xcr isolates as described by Paauw *et al.,* (2024). Briefly, the genomes of 93 *Xanthomonas* strains were sequenced using Oxford Nanopore long-read sequencing and assembled into one-contig whole bacterial chromosome level assemblies. The assemblies were annotated using Prokka v1.14.6 (Seemann, 2014). All predicted ORFs from all assemblies were then clustered into orthogroups using Roary v3.13.0 with default settings (sequence identity threshold >0.95) (Page *et al*., 2015), resulting in a orthogroup presence/absence table. We screened for orthogroups that were present in nearly all Xcc isolates (allowing an orthogroup to be missing in two Xcc genomes to account for potential misassembly or misannotation), and were absent in all Xcr genomes. For this set of 31 orthogroups, we manually inspected the annotation to focus on CWDEs. Using Clinker (Gilchrist & Chooi, 2021), we visualized the genomic flanks of the CWDEs of interest in Xcc8004 and Xcr756c (assemblies from (Paauw *et al*., 2024b))

### Scanning electron microscopy

Scanning electron microscopy was performed as described (Paauw *et al*., 2023). We used four-week-old Savoy cabbage (*B. oleracea* cv. Wirosa) (Vicente *et al*., 2001) plants. To make the sections of the vasculature, the midveins of leaves were dissected by hand with a thin razor blade. Longitudinal cuts along the midvein to reveal the xylem vessels. Samples of approximately 5×5 mm were attached to a sample holder using a thin layer of Tissue-Tek Compound (EMS, Washington, PA, USA). The sample holder was then frozen in liquid nitrogen, and placed into a cryo-preparation chamber (MED 020/VCT 100, Leica, Vienna, Austria). Sublimation of ice on the sample surface was achieved by keeping the sample at −93 °C for approximately 4 minutes. The samples were coated with approximately 12 nm of tungsten, before being loaded into the field emission scanning electron microscope (Magellan 400, FEI, Eindhoven, the Netherlands). The SEM images were taken at −120 °C.

### Leaf disc degradation assay

The leaf disc degradation assay was performed as described (Pfeilmeier *et al*., 2024). Leaf discs of Arabidopsis plants (5.5 weeks old) were collected using a leaf disc puncher (diameter 4 mm) and placed in a 96-well plate filled with 90 µl sterile Milli-Q water. *Xanthomonas* strains were grown for 3 days at 28 °C on KADO agar plates containing 20 µg·mL^-1^ gentamycin. Bacteria were scraped off the plates, resuspended in 10 mM MgSO_4_, washed once, and the bacterial solution was adjusted to OD_600_ of 0.1 in 10 mM MgSO_4_. Leaf discs were inoculated with 10 µl of the inoculum, and were incubated at 21 °C for up to three days. Digital images were taken daily under standardized conditions using a white backlight screen illuminating the plates from beneath. Leaf disc brightness was quantified using a Matlab script (https://github.com/gaebeleinC/leaf-disc_quantification).

### Growth curve assay

Xcc bacteria were grown for 2 days at 28 °C on KADO agar plates containing 20 µg·mL^-1^ gentamycin, before inoculating 10 mL liquid cultures of MOKA medium without antibiotics. These cultures were grown overnight at 28 °C. 2 mL of overnight culture was washed twice, resuspended in M9 medium (M9 salts 1×, glucose 0.4% w/v, MgSO_4_ 1 mM, CaCl_2_ 0.1 mM), and diluted to OD_600_ = 0.5. From this, 15 µl was added to 135 µl medium in microtiter plates, resulting in a start OD_600_ = 0.05. The OD_600_ of the cultures was monitored every 15 minutes for up to 60 hours in a BioTek Synergy H1MF microtiter plate reader.

### Statistical analysis

All statistical calculations were performed in the R programming language 4.2.0. Details on statistical operations are described in the figure legends and in the results section. All boxplots were made with the ‘ggplot2::geom_boxplot()’ function where the middle line represents the median, the upper and lower hinge represent the 75^th^ and 25^th^ percentile respectively, and the upper and lower whisker extend until the largest or smallest value within 1.5 times the interquartile range above the respective hinge. In all boxplots, all individual data points are plotted as points. In experiments with ordinal categorical data or non-normally distributed data, we used a non-parametric Kruskal-Wallis test followed by Dunn’s post-hoc test to detect statistically significant differences between groups. In experiments with continuous, normally distributed data, we used a one-way ANOVA followed by Tukey’s Honest Significant Differences post-hoc test. Groups with no letter in common have a significantly different mean or mean rank (Tukey’s or Dunn’s; α = 0.05).

## Acknowledgements

We would like to thank U. Bonas and D. Büttner for sharing pOGG2. This research was supported by the Topsector T&U program Better Plants for Demands (grant TU18024 to HvdB) and the breeding companies Bejo Zaden B.V. and Rijk Zwaan Breeding B.V.. Both companies were not involved in the design, data analysis, and interpretation of the results.

## Author contributions

**MP:** Conceptualization, Investigation, Visualization, Writing – original draft. **MG**: Investigation. **SP:** Investigation, Writing – review & edit, Supervision. **HvdB:** Conceptualization, Writing – review & edit, Supervision, Project administration, Funding acquisition.

