## Supplementary Material for "Four cell wall-degrading enzymes of *Xanthomonas campestris* pv. *campestris* determine bacterial escape from hydathodes to the leaf vasculature"

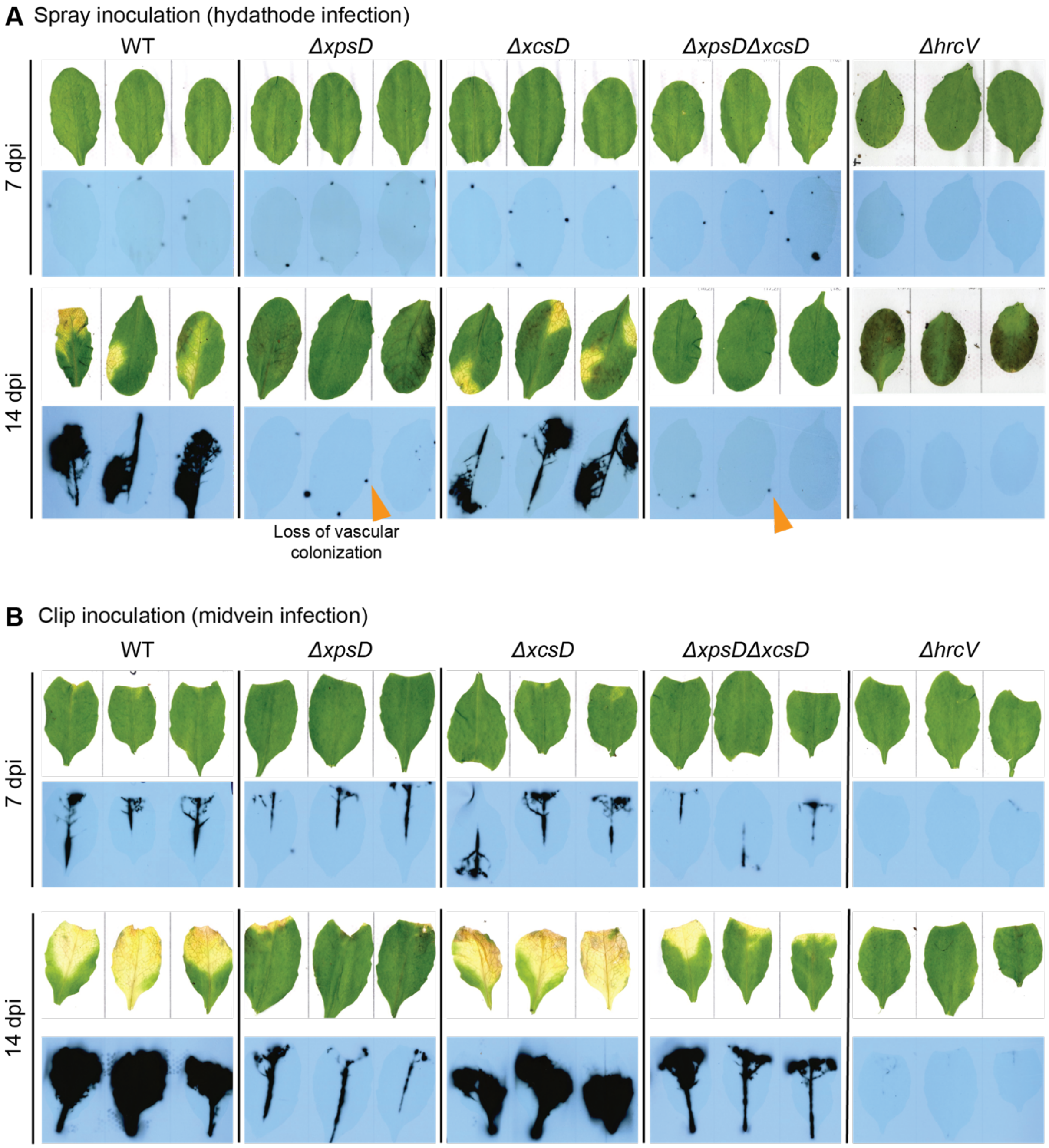


**Figure S1. The Xps type II secretion system is required for Xcc pathogenicity and spread within the host plant in a tissue-specific manner.** Scans of leaves are shown (top row) with the accompanying bacterial bioluminescence signal overlaid on the leaf image (bottom row) at 7 and 14 days post inoculation (dpi) with with Xcc *ΔxopAC Tn7:lux:mTq2* (WT) and indicated mutants by (A) spray inoculation and (B) leaf clipping. Orange arrows highlight infected hydathodes. Subset of images from panel A and B were shown in Figure 1 and 2, respectively.

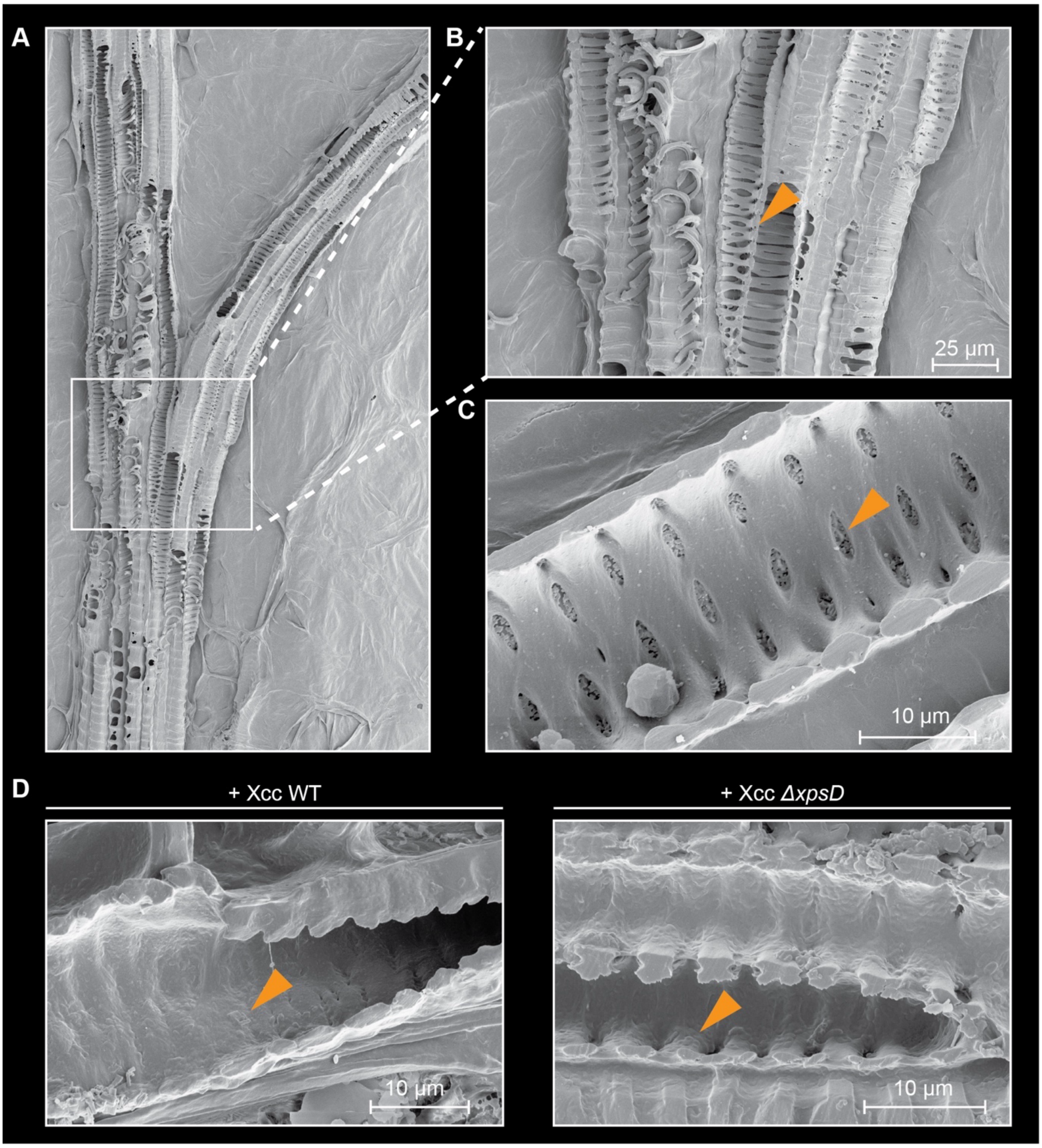


**Figure S2. *Brassica oleracea* xylem structure revealed by scanning electron microscopy.** All panels show longitudinal sections of *B. oleracea* leaves. (A) Overview of a midvein to side vein branch point. White box highlights area zoomed into in panel B. (B) Zoom of highlighted area in panel A. Orange arrow highlights spiraling cell wall structure. (C) Cross-section of individual xylem element. The wall of the xylem element is lined with pits. Orange arrow highlights one pit. (D) Cross-sections of xylem vessels infected with Xcc *ΔxopAC Tn7:lux:mTq2* (WT) or Xcc *ΔxopAC Tn7:lux:mTq2 ΔxpsD*. With both bacterial strains, a biofilm covering the xylem cell walls was observed. A region with bacterial biofilm is highlighted with an orange arrow in both images.

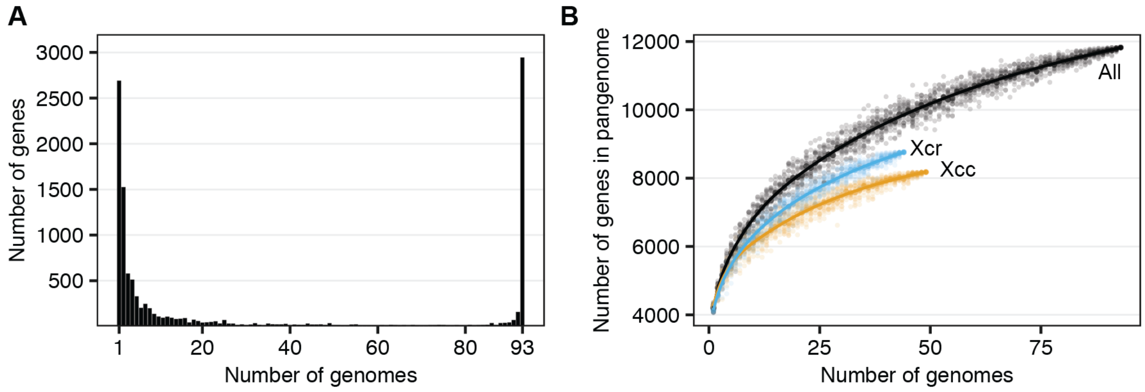


**Figure S3. Xcc/Xcr pangenome statistics.** (A) Gene frequency plot of the 11,824 orthogoups. (B) Rare-faction curves of the total pangenome size for both pathovars together (black), Xcc only (orange) and Xcr only (blue). For each number of genomes, we generated 20 random subset of the genomes and determined pangenome size (individual dots). The line is fitted through the means.

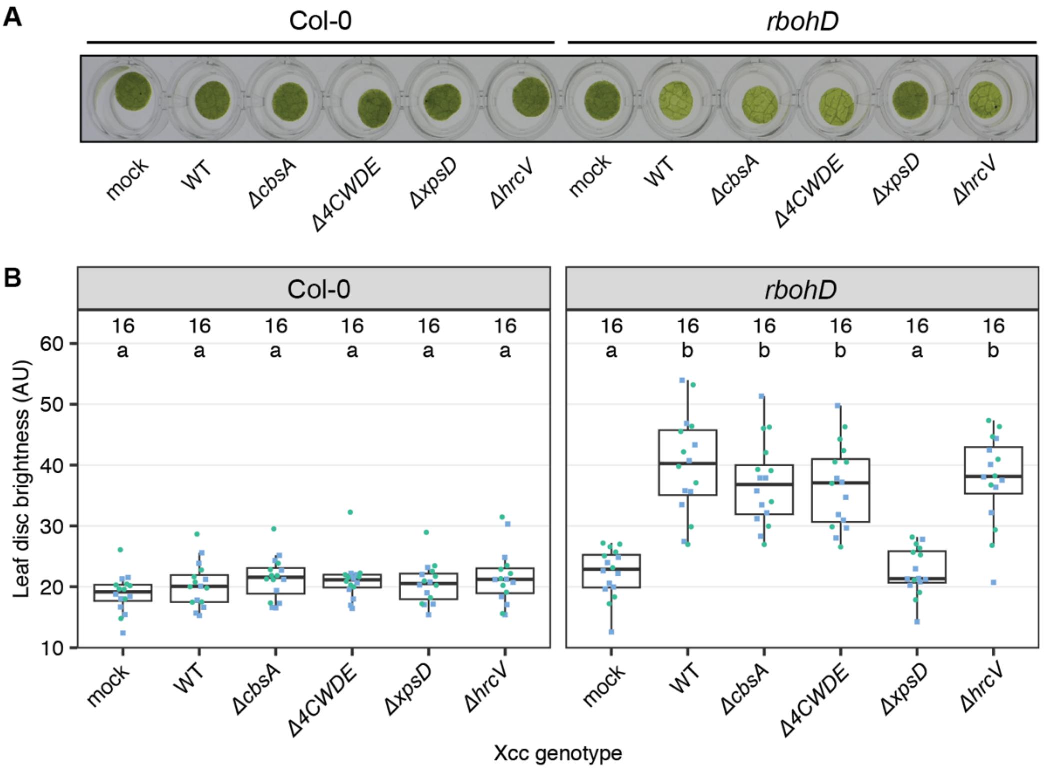


**Figure S4. Degradation of Arabidopsis Col-0 *rbohD* leaf discs is dependent on the Xps-T2SS and not on the four Xcc-specific CWDEs.** (A) Representative pictures of leaf discs, incubated for 40 hours with indicated bacterial cultures. (B) Quantification of the leaf disc brightness of a total of 16 leaf discs for each treatment, divided in two experimental replicates indicated by blue squares or green circles. Treatments with no letter in common have a significantly different mean (Tukey's; α = 0.05). Leaf disc brightness was quantified after 40 hours of incubation.

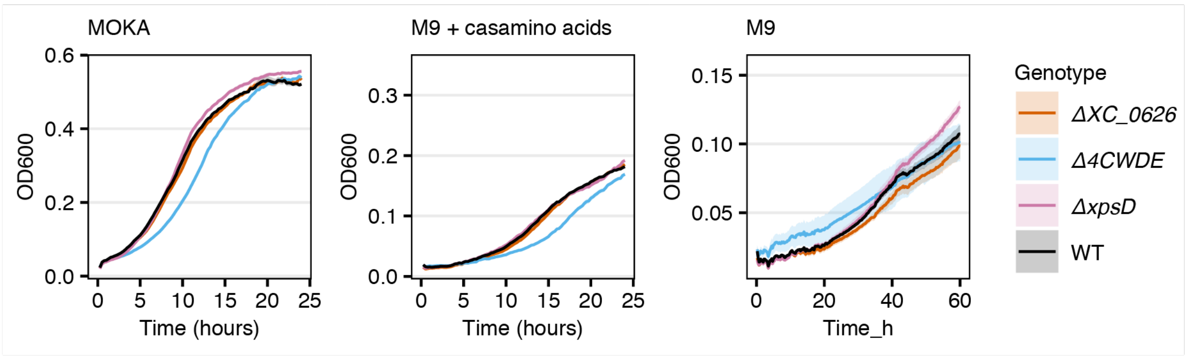


**Figure S5. Growth curves of *Xanthomonas* mutants used in this study.** Bacteria were grown in MOKA medium, minimal M9 medium supplemented with 0.05% casamino acids, or minimal M9 medium. Growth curves were determined in a microtiter plate reader measuring the $OD_{600}$ for 25 or 60 hours.

**Table S1. Orthogroups present in nearly all Xcc isolates, and absent in all Xcr isolates.** For each orthogroup, the Roary orthogroup identifier is indicated, as well as the corresponding gene identifiers in the genome annotations of Xcc8004 from Chapter 4 and two published genome annotations.

| **Roary_group** | **Annotation** | **Paauw *et al.*, 2024** | **Qian *et al.* 2005** (CP000050.1) | **NCBI** (NC_007086.1) |
|---|---|---|---|---|
| group_5781 | hypothetical protein | bc71_00046 | #N/A | XC_RS24260 |
| group_2398 | Saccharopine dehydrogenase NADP binding domain protein | bc71_00047 | XC_0048 | XC_RS00240 |
| group_5783 | hypothetical protein | bc71_00048 | XC_0049 | XC_RS00245 |
| btuB_1 | Vitamin B12 transporter BtuB | bc71_00049 | XC_0050 | XC_RS00250 |
| nagK | N-acetyl-D-glucosamine kinase | bc71_00050 | XC_0051 | XC_RS00255 |
| AvrBs2 | AvrBs2 | bc71_00051 | XC_0052 | XC_RS00260 |
| group_5789 | RNase_H superfamily protein | bc71_00052 | XC_0053 | XC_RS00265 |
| rapA | RNA polymerase-associated protein RapA | bc71_00535 | XC_0536 | XC_RS02725 |
| group_4221 | hypothetical protein | bc71_00536 | XC_0537 | XC_RS02730 |
| group_5809 | Dienelactone hydrolase family protein | bc71_00538 | #N/A | XC_RS24300 |
| glnE | Bifunctional glutamine synthetase adenylyltransferase/adenylyl-removing enzyme | bc71_00559 | XC_0562 | XC_RS02845 |
| XopAG | XopAG | bc71_00560 | #N/A | XC_RS24310 |
| yhhQ | Queuosine precursor transporter | bc71_00561 | XC_0564 | XC_RS02855 |
| group_5832 | endo-β-1,4-glucanase | bc71_00624 | XC_0625 | XC_RS03160 |
| cbhA_1 | Exoglucanase A | bc71_00625 | XC_0626 | XC_RS03165 |
| group_4235 | N-6 DNA Methylase | bc71_01217 | XC_1207 | XC_RS06065 |
| group_4236 | hypothetical protein | bc71_01218 | XC_1208 | XC_RS06070 |
| group_5865 | hypothetical protein | bc71_01219 | XC_1209 | XC_RS06075 |
| group_5866 | hypothetical protein | bc71_01220 | XC_3864 | XC_RS19535 |
| XopK | XopK | bc71_01221 | XC_1210 | XC_RS06080 |
| group_3108 | Secretory lipase | bc71_01755 | XC_1740 | XC_RS08720 |
| fliC | flagellin | bc71_02274 | XC_2245 | XC_RS11300 |
| group_5914 | hypothetical protein | bc71_02892 | XC_2853 | XC_RS14365 |
| group_5917 | Beta-xylosidase | bc71_03199 | XC_3159 | XC_RS15915 |
| XopAM | XopAM | bc71_03200 | XC_3160 | XC_RS15920 |
| XopAY | XopAY | bc71_03216 | XC_3176 | XC_RS16005 |
| XopQ | XopQ | bc71_03217 | XC_3177 | XC_RS16010 |
| group_5920 | hypothetical protein | bc71_03218 | XC_3178 | XC_RS16015 |
| tufA_1 | elongation factor Tu | bc71_03394 | XC_3342 | XC_RS16910 |
| tufA_2 | elongation factor Tu | bc71_03407 | XC_3354 | XC_RS16975 |
| group_5942 | hypothetical protein | bc71_04346 | XC_4276 | XC_RS21660 |

**Table S2. Orthogroups present in nearly all Xcr isolates, and absent in all Xcc isolates.** For each orthogroup, the Roary orthogroup identifier is indicated, as well as the corresponding gene identifier in the genome annotations of Xcr 756c as presented in Paauw *et al.*, 2024 (Chapter 4 of this thesis).

| Roary_group | Annotation | Paauw *et al.*, 2024 |
|---|---|---|
| group_1233 | hypothetical protein | bc01_00049 |
| group_1021 | hypothetical protein | bc01_00050 |
| group_3418 | Saccharopine dehydrogenase NADP binding domain protein | bc01_00051 |
| group_1656 | hypothetical protein | bc01_00052 |
| group_4856 | Autotransporter beta-domain protein | bc01_00828 |
| group_4860 (tufA_2) | elongation factor Tu | bc01_00977 |
| group_4859 (tufA_1) | elongation factor Tu | bc01_00990 |
| group_4862 (rpmD) | 50S ribosomal protein L30 | bc01_01010 |
| group_1713 | Beta-xylosidase | bc01_01186 |
| adhA | putative formaldehyde dehydrogenase AdhA | bc01_01187 |
| group_1714 | Bacterial regulatory proteins, tetR family | bc01_01188 |
| group_4873 | hypothetical protein | bc01_01189 |
| yghA | putative oxidoreductase YghA | bc01_01192 |
| group_3621 | hypothetical protein | bc01_01335 |
| group_3960 | endo-β-1,4-glucanase | bc01_03567 |
| group_3967 (yhhQ) | Queuosine precursor transporter | bc01_03642 |
| group_2283 | putative protein | bc01_03644 |
| group_1554 | Cyclophilin-like family protein | bc01_03646 |
| group_5096 (glnE) | Bifunctional glutamine synthetase adenylyltransferase/adenylyl-removing enzyme | bc01_03647 |
| group_1843 | CE7: acetyl xylan esterase | bc01_03663 |

**Table S3. Bacterial strains used in this study.**

| Xcc genotype | Relevant characteristics | UvA identifier (bglFP) | Reference |
|---|---|---|---|
| *ΔxopAC Tn7:lux:mTq2* | Xcc8004 lacking type III effector XopAC, with chromosomally integrated bioluminescence/fluorescence cassette | 6920 | (Paauw *et al.*, 2023) |
| *ΔxopAC Tn7:lux:mTq2 ΔhrcV* | As bglFP6920, lacking T3SS component HrcV | 8454 | This study |
| *ΔxopAC Tn7:lux:mTq2 ΔxpsD* | As bglFP6920, lacking T2SS component XpsD | 8349 | This study |
| *ΔxopAC Tn7:lux:mTq2 ΔxcsD* | As bglFP6920, lacking T2SS component XcsD | 8351 | This study |

| | | | |
|---|---|---|---|
| *ΔxopAC Tn7:lux:mTq2 ΔxpsD ΔxcsD* | As bglFP6920, lacking T2SS components XpsD and XcsD | 8363 | This study |
| *ΔxopAC Tn7:lux:mTq2 ΔXC_0625* | As bglFP6920, lacking XC_0625 | 8529 | This study |
| *ΔxopAC Tn7:lux:mTq2 ΔXC_0626* | As bglFP6920, lacking CbsA (XC_0626) | 8531 | This study |
| *ΔxopAC Tn7:lux:mTq2 ΔXC_0625 ΔXC_0626* | As bglFP6920, lacking XC_0625 and CbsA (XC_0626) | 8533 | This study |
| *ΔxopAC Tn7:lux:mTq2 ΔXC_1740* | As bglFP6920, lacking XC_1740 | 8535 | This study |
| *ΔxopAC Tn7:lux:mTq2 ΔXC_3159* | As bglFP6920, lacking XC_3159 | 8537 | This study |
| *ΔxopAC Tn7:lux:mTq2 ΔXC_0626 ΔXC_3159* | As bglFP6920, lacking XC_0626 and XC_3159 | 8626 | This study |
| *ΔxopAC Tn7:lux:mTq2 ΔXC_0626 ΔXC_1740* | As bglFP6920, lacking XC_0626 and XC_1740 | 8800 | This study |
| *ΔxopAC Tn7:lux:mTq2 ΔXC_0625 ΔXC_0626 ΔXC_1740* | As bglFP6920, lacking XC_0625, XC_0626 and XC_1740 | 8802 | This study |
| *ΔxopAC Tn7:lux:mTq2 ΔXC_0625 ΔXC_0626 ΔXC_3159* | As bglFP6920, lacking XC_0625, XC_0626 and XC_3159 | 8804 | This study |
| *ΔxopAC Tn7:lux:mTq2 ΔXC_0625 ΔXC_0626 ΔXC_1740 ΔXC_3159* | As bglFP6920, lacking XC_0625, XC_0626, XC_1740 and XC_3159 | 8898 | This study |
| *ΔxopAC Tn7:lux:mTq2 ΔXC_0626* + XC_0626 | As bglFP8531, complemented *in locus* with the coding sequence of XC_0626 | 9002 | This study |

**Table S4. Plasmids used in this study**

| **Plasmid name** | **UvA identifier (pFP)** | **Stored in bacterial glycerol (bglFP)** | **Reference** |
|---|---|---|---|
| pOGG2 | 1398 | N/A | (Solé *et al.*, 2015) |
| pOGG2_XpsD_600 | 2185 | 8339 | This study |
| pOGG2_XcsD_600 | 2188 | 8342 | This study |
| pOGG2_XC_0625 | 2282 | 8461 | This study |
| pOGG2_XC_0626 | 2283 | 8462 | This study |
| pOGG2_XC_0625_XC_0626 | 2284 | 8463 | This study |
| pOGG2_XC_1740 | 2285 | 8464 | This study |
| pOGG2_XC_3159 | 2286 | 8465 | This study |

| | | | |
|---|---|---|---|
| pOGG2_cbsA_KI | 2439 | 8999 | This study |
